# Centromeric variation is shared across ploidy barriers in *Alnus glutinosa* agg

**DOI:** 10.64898/2026.04.21.719804

**Authors:** Jörn F. Gerchen, Bohumil Mandák, Milena Melnyk, Filip Kolář

## Abstract

Polyploidy is often thought to cause immediate reproductive isolation due to sterility and inviability of inter-ploidy offspring. However, recent research demonstrated genome-wide introgression in natural populations of diploids and tetraploids. Yet, it still remains unknown whether introgression varies in strength along the genome and what are the forces underlying such variation. Here we analyzed whole-genome resequencing data from natural populations of the Alder tree, *Alnus glutinosa* agg., which includes a widespread diploid lineage found across large parts of Europe and two autotetraploid lineages with more limited ranges on the Balkan and Iberian Peninsulas. Our sampling involved mixed-ploidy populations, where diploids, triploids and tetraploids co-occur as well as ploidy pure populations. We identified genomic regions of increased admixture, which coincide with putative locations of centromeres. We hypothesize that this pattern of shared variation at pericentromeric regions involves centromere drive, which happens when centromeres increase the likelihood of being included in the oocyte during female meiosis and which has been studied in only a few plant species. While it was previously suggested that centromere drive could cause reproductive isolation and speciation, here we propose that driving centromeres could be able to lift reproductive barriers caused by ploidy differences.

## Introduction

Polyploidy, the presence of multiple genome copies as a result of whole-genome duplication, is common in plants (Otto & Whitton 2000) and genomic analyses showed that polyploidy occurred in several common ancestors of plants (Jiao et al. 2011) and many lineages of animals (Li et al. 2018). One of the main consequences of whole genome duplication is its instant negative effect on reproductive compatibility with diploid relatives (Levin 2002). The offspring of plants with different ploidy levels usually has an uneven number of genome copies, which may either be inviable due to triploid block (Köhler et al. 2010) or sterile due to meiotic issues. As a result, whole-genome duplication has often been considered to cause instant, full reproductive isolation between lineages and thus as a strong driver of speciation. However, as evidence based on modern next generation sequencing technologies showed that gene flow between homoploid species is much more common than previously thought, similar evidence indicating gene flow between lineages with different ploidies is emerging (Monnahan et al. 2019, Bartolić et al. 2025). In combination with older observations form natural populations and crossing results it is becoming apparent that gene flow between lineages with different ploidies is also common (Bartolić et al. 2024) and it has been hypothesized that this gene flow may have contributed to the evolutionary success of polyploid lineages, by providing raw variation that natural selection can act on (Baduel et al. 2018).

Population genomic analyses of genome-wide introgression in homoploid species shows that the distribution of introgressed ancestry can vary substantially across the genome (Martin & Jiggins 2017). Localized peaks of shared ancestry signatures have been identified in many cases and there have been compelling examples which showed adaptive introgression favouring transfer of genetic variation at specific loci (Racimo et al. 2015, Pardo-Diaz et al. 2012, Le Corre et al. 2020, Scott et al. 2025). Other regions retained differentiation between species in the face of strong genome-wide gene flow, a pattern based on so-called speciation loci (Ellegren 2012, Poelstra et al. 2014, Wessing et al. 2023). And a larger scale pattern often showed a positive association between recombination rate and introgression, resulting in relatively lower rates of introgression in regions with reduced recombination (Fu et al. 2022) and in larger chromosomes (Martin et al. 2019).

So far, comparative analyses of introgression patterns across the genome for inter-ploidy introgression, are mostly lacking. Polyploids may differ in the way both purifying selection and positive selection acts on genetic variation due to polysomic masking (Vlček et al. 2025) and it is unclear to what degree these differences can shape genome-wide patterns of introgressed variation. Also gene flow from diploids to polyploids can happen via two specific mechanisms, either via a so-called triploid bridge (Ramsey & Schemske 1998, Husband 2004) or via unreduced gamete formation (Mason & Pires 2015, Kreiner & Husband 2017). A triploid bridge occurs when the triploid offspring resulting from crosses between diploids and tetraploids can form viable haploid or diploid gametes, which can then successfully backcross with diploid or tetraploid relatives, respectively (Ramsey & Schemske 1998). Depending on the proportion of viable haploid and diploid gametes produced by triploids, this is expected to allow bi-directional gene flow. Alternatively, diploids can produce a fraction of unreduced, diploid, gametes, which can then cross with tetraploids leading to tetraploid F1 hybrids, allowing for unidirectional gene flow from diploids to tetraploids (Ramsey & Schemske 1998, Brown et al. 2024). So far it is unclear if these two pathways for inter-ploidy gene flow could specifically shape the patterns of introgressed variation. This could for example be the case if specific structural or genetic variants favour the production of unreduced gametes or increase the fertility of triploids. In addition, in both diploids and polyploids genomic regions of low recombination may not always behave based on the assumptions of Mendelian segregation, which underlie the population genetic assumptions for gene flow. This may be true for centromeres, which are commonly surrounded by large non-recombining regions and which may show deviations from mendelian segregation due to centromere drive.

Centromere drive causes deviations from Mendelian segregation by selfish centromeres, which favour their own transmission during female meiosis (Dudka et al. 2022). This happens because only one of four meiotic products will end up in the oocyte nucleus and driving centromeres increase their likelihood of being included in the oocyte during meiosis 1 (Kursel et al. 2018). This process is often associated with structural variation at the centromeres and differences in the composition of centromeric repeats (Akera et al. 2019). Centromere drive is also thought to be one factor underlying the so-called centromere paradox, which is based on the observation that centromeres are among the most variable regions in the genome, while the meiotic machinery that interacts with it is highly conserved (Henikoff et al. 2001). This high variability at centromeres could be driven by repeated rounds of completed centromeric drive, i.e. newly formed driving centromeres getting fixed in populations (Henikoff et al. 2001). Centromere drive has been hypothesized to play a substantial role in reproductive isolation (Searle & Pardo-Manuel de Villena 2024). However, it is conceivable that driving centromeres could also be particularly effective at crossing strong reproductive barriers specifically because their transmission advantage may outweigh negative fitness effects of admixed variation in hybrids. Other types of meiotic drive elements have been shown to cross species barriers (Meiklejohn et al. 2018, Svedberg et al. 2021), but this has not been observed for driving centromeres in plants. One reason may be that driving centromeres tend to fix quickly and leave few detectable population genomic signatures. However, inter-ploidy introgression could retain such population genomic signatures by slowing the spread of driving centromeres in the population, due to the higher copy number of polyploid genome copies and due to polysomic masking of linked deleterious variation, which would be removed by purifying selection in diploids. Here we propose that population genomic evidence from the Alder tree, Alnus glutinosa agg. could be consistent with such a scenario.

The Alder tree, *Alnus glutinosa* agg. (Betulaceae) is a common wetland tree found in large parts of Europe, Northern Africa and southwest Asia which shows variation in ploidy (Vít et al. 2017). There is one widespread diploid lineage (*A. glutinosa*), which is found across large parts of Europe and two autotetraploid lineages likely representing two independent whole genome duplication (WGD) events, *A. rohlenae* and *A. lusitanica*, found on the Balkans and on the Iberian Peninsula, respectively (Mandák et al 2016). Triploids were found in contact zones between diploid *A. glutinosa* and tetraploid *A. rohlenae* (Šmíd et al. 2022), but so far no triploids were found in a large survey of cytotypes on the Iberian peninsula (Cuerdo et al. 2025). Population genetic evidence suggested some evidence of interploidy admixture between diploid *A. glutinosa* and tetraploid *A. lusitanica* (Martin et al. 2024), but so far a comprehensive analysis of local patterns of interploidy introgression within a genome using population genomic data was lacking in both species. Here we present a population genomic analysis, which combines whole genome re-sequencing covering the range of all three lineages, including a large newly discovered contact zone between *A. lusitanica* and *A. glutinosa* in northern Spain, where we found large numbers of triploids. Our population genomic analysis suggests low rates of gene flow for most of the genome, but it also shows extensive interploidy sharing of haplotypes at putative centromeric regions. We propose that this could be the result of driving centromeres crossing ploidy barriers at multiple chromosomes and in both contact zones.

## Material and Methods

Our sampling consisted of samples that were collected for a previous publication (Šmíd et al. 2022) as well as new samples (Suppl.Table 1). Leaf material was collected from trees at least 10 m apart and dried on silica gel. Genomic DNA was extracted using the sorbitol extraction method (Štorchová et al. 2000) and then purified using AMPure XP (Beckman Coulter Inc., Brea, California, USA). We resequenced a total of 136 samples, which included *A. glutinosa*, *A. lusitanica* and *A. rohlenae* from the Balkans and the Iberian peninsula as well as one sample of *A. incana* and one of *A. orientalis*, which we used as outgroup samples.

### Genotyping

We generated Illumina short read sequencing libraries using the LITE (Perez-Sepulveda 2021) protocol and samples were sequenced on the Illumina NovaSeq platform. We genotyped the *Alnus* samples using our self-developed mixed-ploidy variant calling pipeline (https://github.com/jgerchen/polyploid_variant_calling) by aligning Illumina reads against the *A. glutinosa* genome assembly developed by the Darwin Tree of Life Project (Christenhusz et al. 2024). For this workflow we first trimmed paired-end reads using trimmomatic 0.39 (Bolder et al. 2014) with standard parameters and aligned reads against the reference genome using BWA mem 0.7.18 (Li, 2013) and we sorted aligned reads by position using samtools 1.20 (Danecek et al. 2021). After merging multiple libraries for the same sample using samtools merge we removed PCR duplicates using the MarkDuplicates comand from PicardTools 3.1.1 (Broad Institute, 2019). We then used GATK HaplotypeCaller 4.6.20 (Poplin et al. 2017) for combined genotyping. For this we first ran HaplotypeCaller per sample with the options —minimum-mapping-quality and —base-quality-score both set to 20 and the option –ERC set to GVCF to generate a GVCF output file. We then used the GenomicsDBImport module to build a combined Genomics DB for all samples and ran combined genotyping using the GenotypeGVCFs module with the option --include-non-variant-sites activated so that we genotyped both variant and invariant sites.

Our filtering of genotypes involved a newly developed machine learning approach for distinguishing true positive biallelic SNPs from sequencing errors in non-model species like *A. glutinosa* for which a reference panel of SNPs is lacking. The idea behind this approach was that sequencing errors typically occur at low frequencies in the population and variants that are found in multiple individuals are likely to be true positives. Based on this we selected a subset of SNPs that had alt alleles in multiple (four or more) individuals, and modified copies of the bam files generated in the earlier steps of our variant pipeline to turn them into low frequency variants by changing the nucleotides at the variant positions from alternative to reference allele in a subset of samples. We then ran a separate round of genotyping for all samples and trained a machine learning model on six INFO statistics (FS, QD, MQ, MQRankSum, ReadPosRankSum, SOR) from the resulting raw VCF file based on modified sites as well as on a number of unmodified sites relative to the site frequencies based on the original data. We then applied this model to the original data to get an error probability for all biallelic SNPs.

This approach used four custom python scripts for each step of the machine learning approach. We generated our training data based on a subset of two chromosomes (Chr3 and Chr4) to reduce computational overhead. With the first script, training_select_variants.py, we selected a set of 50000 variants, which have to be at least 1kb apart and which must have alt alleles in at least four individuals and uses the numpy 1.26.4 (Harris et al. 2020) and matplotlib 3.10.7 (Hunter, 2007) python modules. The second script, replace_variants.py, then uses the list of variants generated by the previous script to modify copies of bam files for each sample. It uses pysam 0.23.3 (https://github.com/pysam-developers/pysam) to modify bam files and the resulting modified bam files were resorted using samtools 1.23 (Danecek et al. 2021). Then genotyping on the modified bam files was done using GATK HaplotypeCaller with the same parameters as described above, with the only exception that we did not set the --include-non-variant-sites for the GenotypeGVCFs module, because here we only needed variants. The resulting VCF file was used to train a machine learning model in the script pu_learn.py. This script uses the pulearn 0.0.11 python package (https://pulearn.github.io/pulearn/), which implements a positive-unlabeled approach (Elka & Noto 2008) for training a classifier model, which distinguished our true positives (sites from the previous steps) from a mixture of true and false positives (all other biallelic SNPs in the dataset). The underlying classifier model is the HistGradientBoosting classifier implemented in scikit-learn 1.5.2 (Pedregosa et al. 2011). The fourth script, pu_predict.py, applies the classification model generated in the previous script to all biallelic SNPs from the original VCF file and infers the probability of being included in the truth set, PUprob, for each biallelic SNP. This script also generated a plot of ts/tv values of biallelic SNPs sorted by PUprob. We used this plot to manually determine a filtering cutoff by finding a PUprob value, which cuts off the peak of ts/tv values at the left side of the plot.

For filtering of variants we first used the annotate module of bcftools to add PUprob values for biallelic SNPs. We separated biallelic SNPs and invariant sites and removed all indels, complex variants and multiallelic sites. We retained only biallelic SNPs with a PUprob value greater than our previously set value, merged the VCF file with the invariant sites again and used the filter module of bcftools to set all genotypes with a sequencing depth smaller than 4 to no-call. We then used the +fill-AN-AC module to update values for allele counts and allele number and the –a option of bcftools view to remove alt alleles, which were not present in the file after the previous filtering steps. Finally, we used a custom python script (https://github.com/jgerchen/polyploid_popgen/tree/main/depth_mask) on the filtered VCF file to identify sites with disproportionately high sequencing depth. This script identifies sites where more than half of samples have a sequencing depth greater than the mean depth of the sample plus two times its standard deviation. We then used bcftools view to exclude high depth sites and create our final VCF file.

### Population genomic analysis

We generated a pruned VCF file for several of the population genomic analyses. We used the prune module of bcftools with a window size 100 kb, a maximum of 10 sites per window and a maximum LD of 0.2. We then removed retained sites with a minor allele frequency smaller than 0.05 using bcftools view.

We used the pruned VCF file to generate a NeighborNet using a custom R script (https://github.com/jgerchen/polyintro/blob/main/workflow/scripts/nei/neis_distance.r) in R 4.5 (R Core Team 2025), which uses the R packages StAMPP 1.6.3 (Pembleton et al. 2013), vcfR 1.15 (Knaus et al. 2017), adegenet 2.1.11 (Jombart et al. 2011) and phangorn 2.12.1 (Schliep et al. 2017).

We used the pruned VCF file to generate an input file for Structure using a custom python script (https://github.com/jgerchen/polyintro/blob/main/workflow/scripts/structure/ vcf_to_structure.py) and ran 10 replicates of STRUCTURE 2.3.4 (Pritchard et al. 2000) for each of k=1-7 clusters with a burn-in length of 10,000 and data collection for 50,000 iterations. We then used CLUMPP 1.1.2 with the M option set to 2 and the greedy option set to 2 and the number of permutation repeats to 10,000 (Jakobsson et al. 2007) to generate average values for each value of k.

### Estimation of D statistics

We used scripts by Simon Martin (https://github.com/simonhmartin/genomics_general) to estimate D statistics in 50kb sliding windows for each of the three contact zones. For each contact zone we set a tetraploid population, which is distant from the contact zone as P1 population, tetraploids from within the contact zone as P2 population, and diploids from either within the contact zone or from a closely located pure diploid population as P3 population. In all three cases we used the *A. incana* sample and the *A. orientalis* sample as outgroup.

### Local PCAs

We used lostruct (Li & Ralph 2019) to test how local population structure differs between centromeres and chromosome arms. We ran local PCAs separately for the three different mixed-ploidy populations and we included all diploids, triploids and tetraploids from each mixed-ploidy population as well as one more distant tetraploid population and one close diploid from the same region. We estimated local PCAs in 50kb windows with k=2 and estimated the pairwise distances between PCAs in windows using the pc_dist function of lostruct. We then ran a multidimensional scaling analysis on the pairwise distances using the cmdscale function in R with four dimensions. We plotted the MDS values of the four dimensions separately for each contact zone and identified representative PCAs for the chromosome arms and outlier windows for the centromeres. Depending on the MDS plots, we selected either maximum or minimum values on the first MDS dimension to identify windows, with PCAs, which are representative for population structure on the chromosome arms. For the centromeres, we identified the MDS dimension, which had the strongest outlier peaks in either direction and selected the window with the most extreme MDS value for that peak. In addition, we also chose PCA windows overlapping with the window with the most extreme D-statistics for one of the chromosomes with D-statistics peaks for each of the contact zones.

### Neighbor-joining trees

We developed a custom R script, which is based on R 4.5 (R Core Team 2025) and which uses vcfR 1.15 (Knaus et al. 2017) and adegenet 2.1.11 (Jombart et al. 2011) to read the VCF file and split the it into 100kb sliding windows. We then estimated pairwise distances using Nei’s D estimated by StAMPP 1.6.3 (Pembleton et al. 2013) and build neighbor-joining trees using the nj function in ape 5.8-1 (Paradis et al. 2004). We plotted local neighbor-joining trees overlapping with the window with the most extreme D-statistics for one of the chromosomes with D-statistics peaks for each of the contact zones.

### Identification of centromeres

We used two approaches with the aim to identify centromeres and pericentromeric regions based on their repeat content. First, we ran RepeatModeler 2.0.4 (Flynn et al. 2020) to identify a de-novo library of transposable elements from the *Alnus glutinosa* reference genome and we used the RepeatMasker 4.1.5 (Smit et al. 2013-2015) to mask repeats. Then we ran CentiER 3.0 (Xu et al. 2023), which identifies putative centromeric regions based on both tandem repeats and TE content.

## Results

### Population structure

Our population genomic analyses showed that diploids and tetraploids form distinct groups. Our NeighborNet forms three major clades representing diploids, tetraploids from the Balkans and tetraploids from Iberia (Fig. 1A). While the diploids from the Balkans and Iberia form a single clade, the two tetraploid lineages form two clearly diverged clades, suggesting either independent origins or substantial historical divergence. Similarly, while our Structure results for k=2 clusters separates tetraploids from Iberia from diploids and tetraploids from the Balkans, results for k=3 further separates tetraploids based on Balkans and Iberian origins and results for k=4 indicate that there may be additional variation within tetraploids from the Balkans (Fig. 1C).

**Fig 1:**
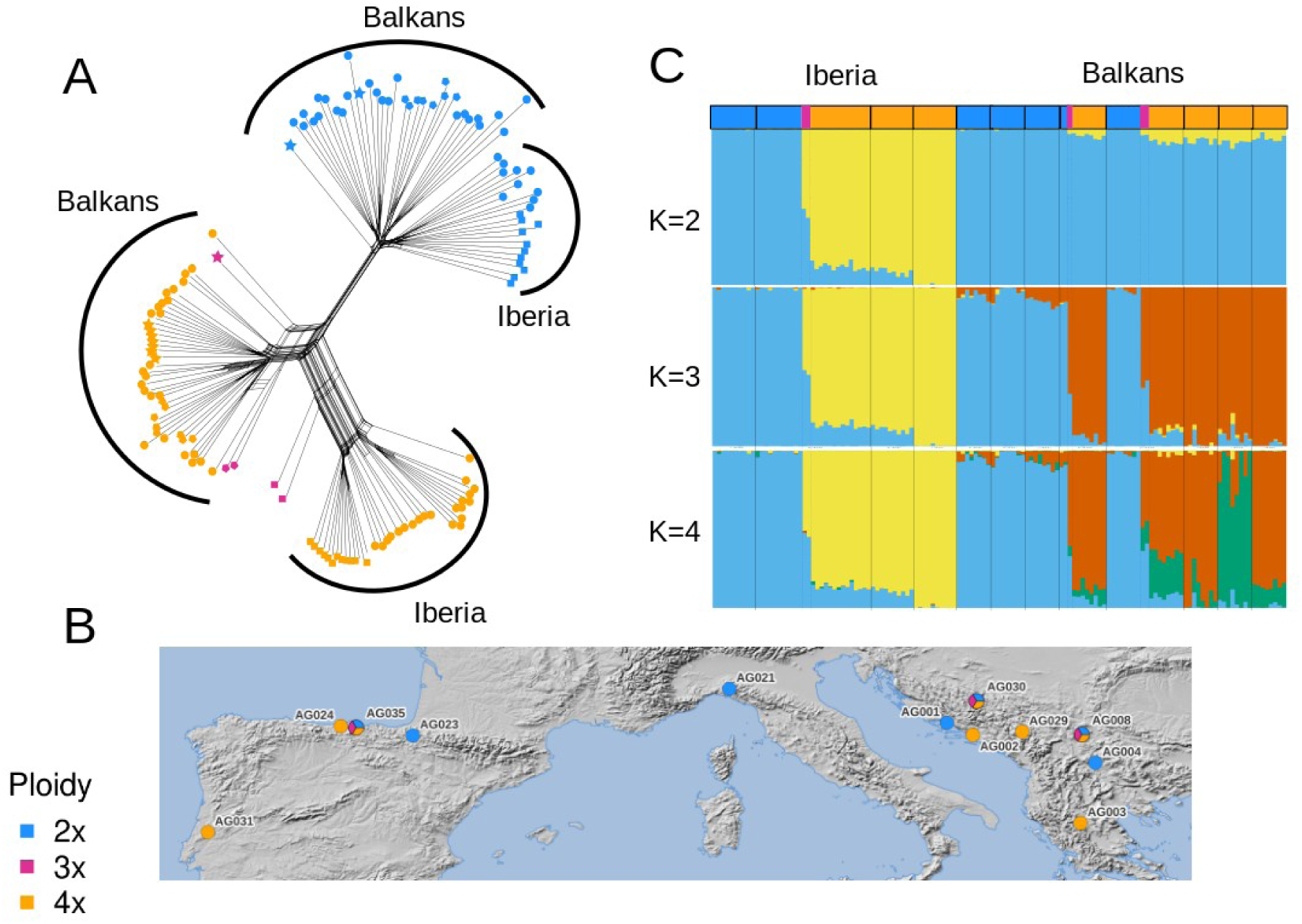
Population structure and sampling of *A. glutinosa* agg. populations on the Balkans and the Iberian peninsula based on 33272 genome-wide SNPs. A: NeighborNet of individual samples. B: sampling of *A. glutinosa* populations. Locations with multiple colors indicate mixed-ploidy populations C: Results of STRUCTURE analysis for K values from 2 to 4.

### Repeat content and centromeres

We identified single peaks for regions of elevated repeat content at the center of each chromosome (Fig. 2). These regions corresponded with the putative locations of pericentromeric regions identified by Centier.

**Fig. 2:**
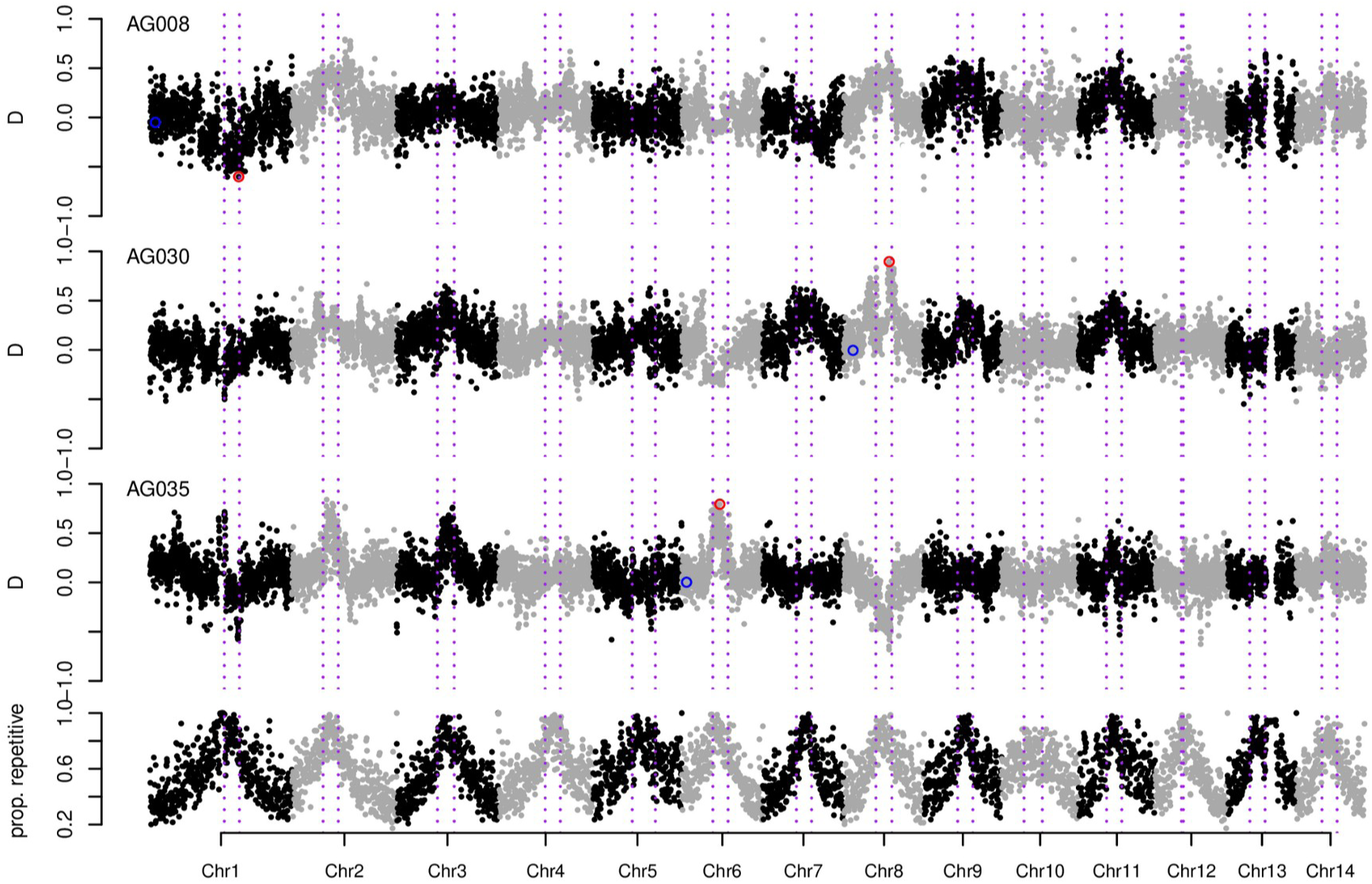
D-statistics and repeat content for three mixed-ploidy populations of *A. glutinosa*. The three top rows show mean D-statistics in 50kb windows for each contact zone, with positive D indicating an excess of diploid-like ancestry in tetraploids from the contact zone and negative D-statistics indicating an excess of diploid-like ancestry in the tetraploid population from outside the contact zone. Red and blue circles indicate windows used for PCAs and Neighbor-joining trees at centromeres and chromosome arms plotted in Fig. 3. The bottom row indicates the proportion of repetitive sequences in 100kb windows and the dotted purple lines indicate the putative pericentromeric regions inferred by Centier.

### Gene flow

Our analyses of D-statistics for the three contact zones showed substantial variation across the genome. Outlier values of D-statistics were concentrated at the center of chromosomes and were located in or close to regions, which we also identified as putative pericentromeric regions using Centier (Fig. 2). Notably, these outliers included both positive (indicating an excess of shared variation between tetraploids from the contact zone and diploids, for example on Chr2 and Chr8 for contact zone AG008, Chr3 and Chr8 for contact zone AG008 and Chr2 and Chr6 for contact zone AG035) and negative D statistics (indicating an excess of shared variation between tetraploids from outside the contact zone and diploids, for example Chr1 for contact zone AG008 and Chr8 for contact zone AG035).

### Local PCAs

All three of our genome-wide local PCA analyses showed varying peaks across dimensions 2, 3 and 4 of our MDS analyses. All of these peaks were located at the centers of chromosomes (Fig. S2, S5 and S8), and correspond with the peaks for D-statistics, which were also localized in the centromeric regions (Fig. 2). Importantly, MDS peaks at different chromosomes in the same mixed-ploidy population were found on different MDS dimensions, and some peaks were going in different directions. This indicates that the identities of outlier individuals that cause these peaks vary between chromosomes. In contrast, patterns for MDS1 were more predictable in all three mixed-ploidy populations with outlier windows located on all chromosome arms. The local PCAs on chromosome arms generally showed a pattern, which clearly separates diploids and tetraploids from both within and outside the contact zone along PCA axis 1, while triploids were located either at an intermediate position or closer to tetraploids (Fig. S4, S7 and S10). In contrast, PCAs from windows corresponding to outlier peaks showed substantial variation in the relative locations of diploids and tetraploids. For most outlier PCAs diploids formed a single cluster, while a varying proportion of tetraploids from either within or outside the contact zone were co-localized with the diploids, while the others formed one or multiple separate clusters along PC1 (Fig. S3, S6 and S9). This pattern would be consistent with some proportion of tetraploids containing diploid-like variation at centromeric regions. The presence of multiple discrete tetraploid clusters separated at PC1 (For example Chr2 for contact zone AG008, Chr4 for AG030 and Chr9 for AG035) could be caused by variable copy numbers of diploid-like haplotypes in tetraploids. However, in some cases (Chr1 for AG008, Chr13 for AG030 and Chr1, Chr7 and Chr14 for AG035) also multiple separate diploid clusters, were found which could indicate that a small number of diploids also have otherwise predominantly tetraploid-like variation at the centromeres.

The PCA windows we selected based on D-statistics instead of MDS analyses were consistent with the pattern observed for MDS outliers (Fig. 3). PCA windows from chromosome arms with D-statistics close to zero showed a clear separation between diploids and tetraploids and the corresponding neighbor-joining trees separated diploids and tetraploids in two separate clades. In contrast, PCA windows corresponding to centromeric D-statistics outliers showed similar patterns as the outlier windows from the MDS analyses, with some tetraploids clustering with diploids. Here the corresponding neighbor-joining trees did not form two clearly separate clades for diploids and tetraploids, but diploids and tetraploids were intermixed to varying degrees (Fig. 3).

**Fig. 3:**
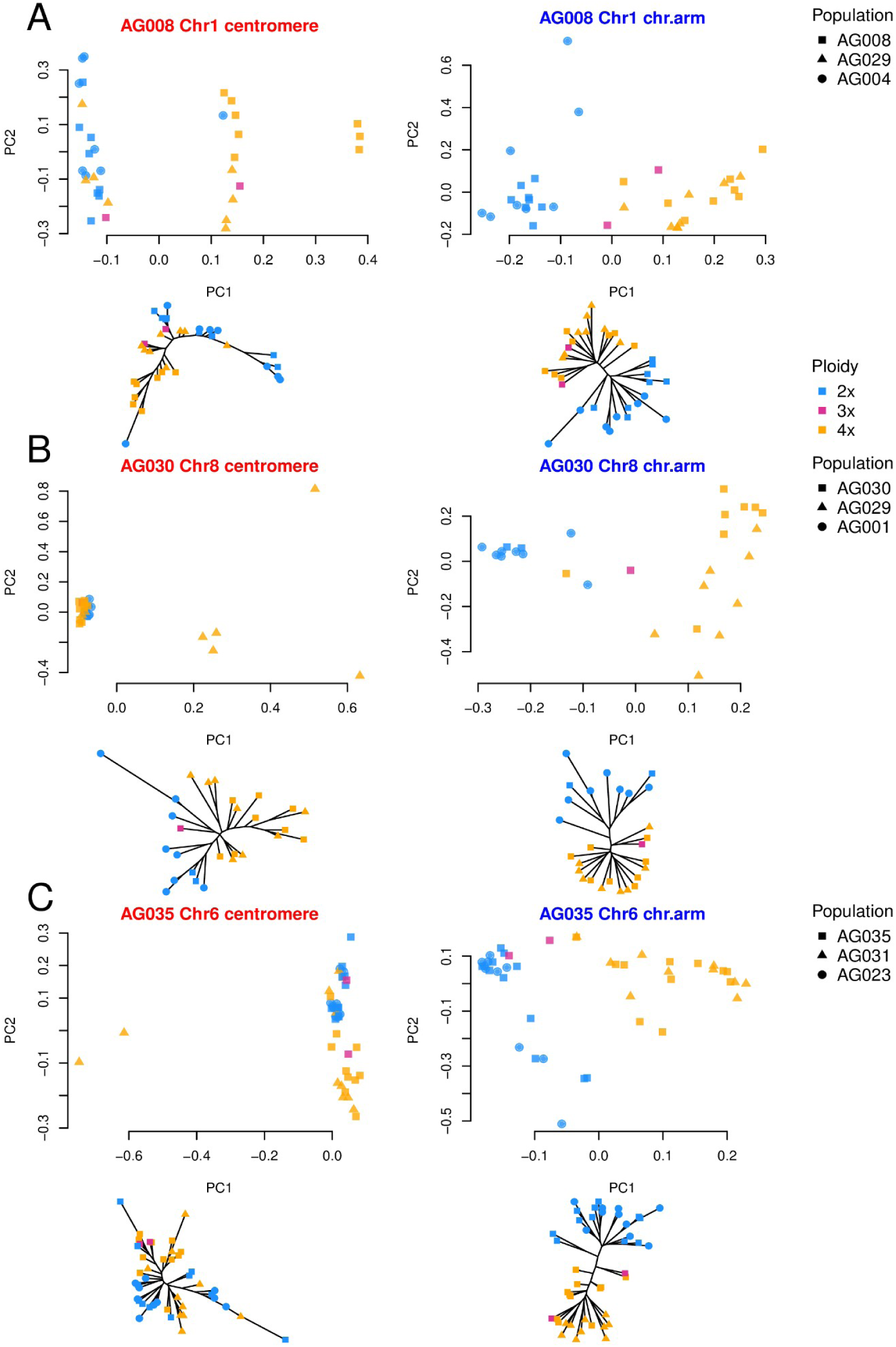
Local PCAs and neighbor-joining trees for example chromosome arms and from centromeric regions for the three mixed-ploidy populations.

## Discussion

Here we provide the first in depth population genomic analysis of diploid and tetraploid lineages of *A. glutinosa* agg. In addition, we identified a new large contact zone between diploid *A. glutinosa* and tetraploid *A. lusitanica*, where diploids and tetraploids co-occur for several kilometers along the river Asón in northern Spain and several of its smaller tributaries up to its estuary into the Atlantic (Fig. S1). Unlike in previous studies of *A. glutinosa* and *A. lusitanica* on the Iberian peninsula (Martín et al. 2024, Cuerdo et al. 2025) we also identified adult triploid trees, suggesting ample opportunities for hybridization and inter-ploidy gene flow via a triploid bridge.

Our genome-wide population genomic analysis suggests a clear separation between ploidies within each contact zone and between tetraploid lineages from the Balkans and Iberia (Fig. 1). These results are in stark contrast to the results from our D-statistics analyses on a local genomic scale, where we tested for the presence of shared genetic variation between diploids and tetraploids from the contact zones relative to tetraploid populations from outside the contact zone. Here we identified substantial shared genetic variation between diploid and tetraploid populations of *A. glutinosa* in both mixed-ploidy populations from the Balkans (AG008 and AG030) as well as at the mixed-ploidy population from Iberia (AG035) (Fig. 2). Importantly, the strongest outliers of D-statistics are localized in putative pericentromeric regions identified by CentiER and which also coincided with peaks of repetitive sequence content. We identified peaks of excessive D-statistics in these regions, suggesting variation in the extent of diploid-like variation found in tetraploid populations.

Our individual-based local PCAs indicated that the difference between D-statistics were also reflected in the distribution of individual genetic variation on chromosome arms and centromeres. On chromosome arms, where local estimates of D-statistics were on average close to zero, most local PCAs clearly separated diploid and tetraploid populations (Fig. S4, S7, S10). In contrast, in pericentromeric regions, which were the locations where both outliers of D-statistics and peaks of MDS analysis of principal components were found, variable numbers of tetraploid individuals clustered close to diploids while other tetraploids remained separate (Fig. S5, S8, S11).

Taken together, these results imply that there are large pericentromeric regions, where diploid-like genetic variation is shared across ploidy barriers. The strong difference between chromosome arms and pericentromeric regions and the repeated observation of peaks of shared variation across centromeres and contact zones suggests that these patterns could be related to specific properties of pericentromeric regions, which allow the ancestrally diploid variation to preferentially introgress across ploidy barriers. Here we suggest that centromere drive, the preferential transmission of centromeres during female meiosis (Dudka et al. 2022), could cause the observed patterns of shared genetic variation across ploidy barriers. Importantly, although the identity of chromosomes differed, this general pattern of excessive sharing of mostly diploid-like pericentromeric regions in tetraploid populations was found in all three mixed-ploidy populations (Fig. 2), thus affecting two diverged tetraploid lineages, *A. rohlenae* on the Balkans and *A. lusitanica* on the Iberian peninsula.

We consider that centromere drive could cause these observations, because it specifically affects the centromeres and because its non-Mendelian inheritance could favour spread of introgressed regions beyond the mixed-ploidy populations, similar to observations in other meiotic drive systems (Meiklejohn et al. 2018, Svedberg et al. 2021). Our hypothesis is based on what we know about how centromere drive works in other organisms, where it can be caused by structural variation, like the extension of centromeric repeats (Akera et al. 2019). Such variation can evolve rapidly via the interplay of recombination, which homogenizes satellite repeats, and that of transposable element insertion and mutation, which disrupts homogeneous repeats (Malik & Hernikoff 2009). The rate of emergence of such variation could be affected by differences in demography and evolutionary history of diploids and tetraploids. The more widespread diploid *A. glutinosa* lineage is expected to have a larger effective population size, and driving centromeres could arise at a higher rate than in tetraploids. As a result, diploids may have accumulated a set of stronger driving centromeres relative to tetraploids. This could explain the largely unidirectional pattern where diploid-like shared variation is found in tetraploids, while tetraploid-like variation is found in diploids in only a few cases.

Furthermore, genes involved in counteracting centromere drive in plants, in particular CENH3 which encodes centromeric nucleosomes, interact with the kinetochore complex (Finseth et al. 2015, Finseth et al. 2021). Several other genes that encode core components of the kinetochore complex have been shown to be involved in adaptation to whole-genome duplication (Bray et al. 2024). This raises the intriguing possibility that there could be an interaction between adaptation to whole-genome duplication and susceptibility to driving centromeres from diploid relatives. If driving centromeres from diploids could affect meiotic stability in tetraploids, because they are not co-adapted to tetraploid meiosis, balancing selection could prevent fixation of such diploid-like driving centromeres in tetraploids and vice versa. This could result in a mosaic of centromere types across the range of tetraploids, as we observe here, and would be comparable to centromere drive in *Mimulus gutatus*, where reduced male fitness prevents fixation of a driving allele (Fishman et al. 2008).

Our results are based on aligning short read sequencing data against centromeric regions of the *A. glutinosa* genome assembly (Christenhusz et al. 2024), which have a strong enrichment of repetitive sequences like transposable elements and centromeric repeats (Fig. 2). Aligning short reads against such regions can result in substantial biases and introduce different types of artifacts like disproportionally high sequencing coverage and false variant calls. While our data are unlikely to be free from these issues, we believe that the general patterns of shared variation between diploids and tetraploids at the centromeres we observed here do not present artifacts, but are based on real sequence differences for multiple reasons. First, the observed patterns span large, megabase-scale regions (Fig. 2), which are much larger than the genomic regions covered by individual Illumina short read pairs. To introduce significant amounts of spurious variation, sequencing artifacts would have to affect thousands of reads in the same region in the same way, which seems unlikely. Second, there is substantial variation in the distribution of alternative genotypes between chromosomes, individuals, populations, contact zones and lineages (Fig. 2, S2, S5 and S8). This observation is at odds with systematic errors like alignment artifacts, which would be expected to consistently affect individuals when aligned to the same reference genome.

Some of our population genomic analyses may be biased when comparing samples of different ploidies. PCA has been shown to be biased in mixed-ploidy comparisons, resulting in a greater spread tetraploid than diploid data, especially in cases when there is low differentiation between ploidies (Meirmans et al. 2018). This may also cause some notable differences in the shape of PCA analyses and the topology of neighbor-joining trees (Fig. 3B, Fig. 3C), however both trees and PCAs from chromosome arms clearly show strong differentiation between ploidies, while those from centromeres do support a varying degree of admixture (Fig. S2-S10). The results of PCA and distance-based metrics like Nei’s D may also give inconsistent results if individuals have different numbers of diverged (tetraploid-like) haplotypes. Further analyses using long read data will likely help us to understand how copy number variation and structural variation is involved in the observed patterns, since they would allow us to both properly phase haplotypes and to detect structural variation, which remains undetectable using our current short read data.

## Conclusions

Our population genomic dataset on diploid and tetraploid *A. glutinosa* shows an intriguing pattern, where an excess of diploid-like variation is found in tetraploids of *A. glutinosa* in putative pericentromeric regions, suggesting increased admixture at the centromeres. We find these results in three mixed-ploidy populations in two diverged tetraploid lineages and we hypothesize that it could be caused by centromere drive crossing ploidy barriers. Further analysis using long read sequencing and potentially experimental evidence could further test this hypothesis. These results have the potential to show a previously unconsidered route how variation may cross species barriers and it could provide additional examples of centromere drive in plants.

## Acknowledgements

The authors thank J. Vítová and J. Nosková for preparation of sequencing libraries. This work was funded by a Postdoc Individual fellowship of the Czech Science Foundation (GAČR) with the grant number 23-07275I and The European Research Council (ERC) under the European Union’s Horizon 2020 research and innovation programme (ERC-StG 850852 DOUBLE ADAPT led by F.K.). Computational resources were provided by the e-INFRA CZ project (ID:90254), supported by the Ministry of Education, Youth and Sports of the Czech Republic.

## Supplementary Information

**Fig. S1:**
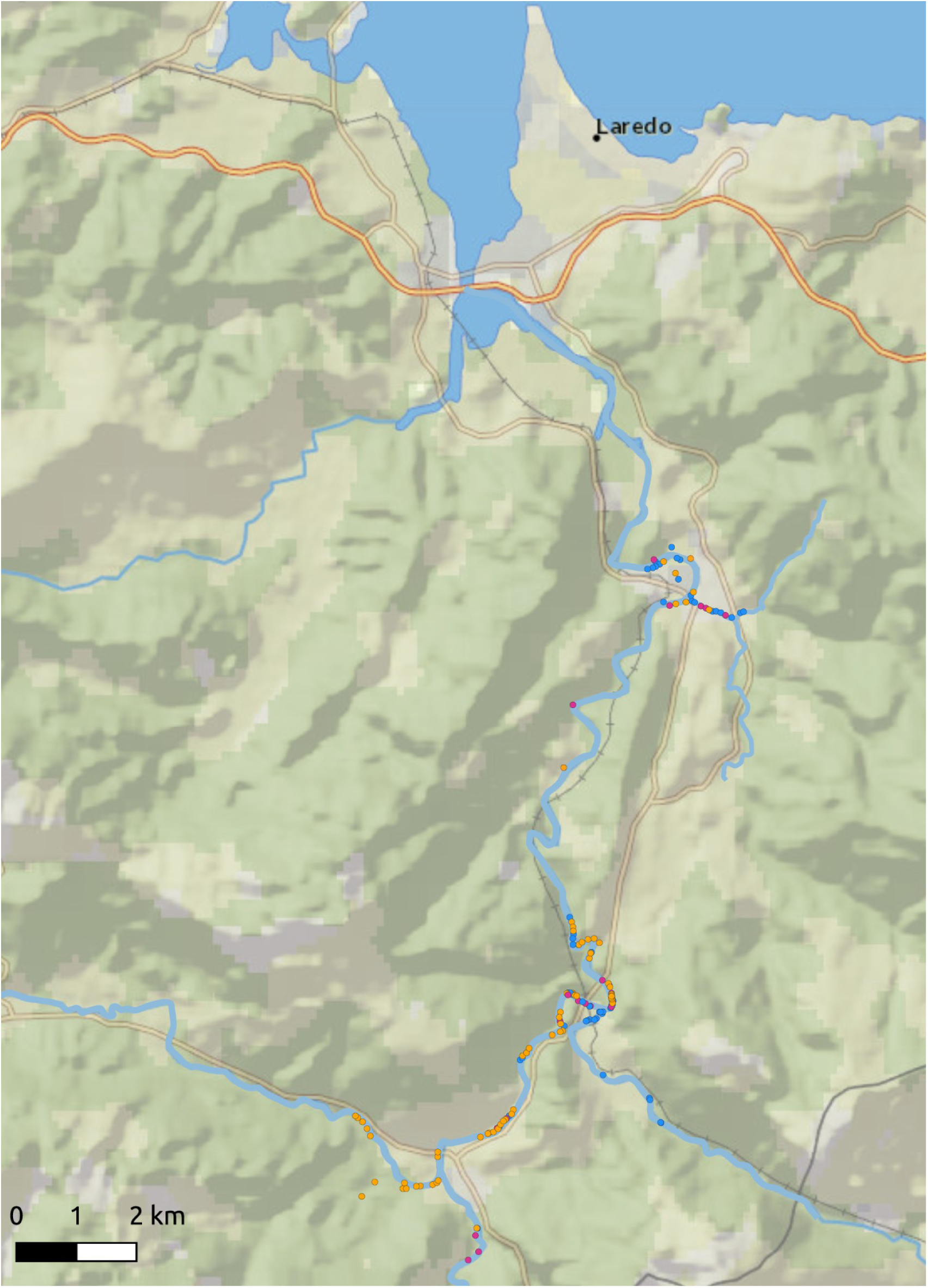
Sampling map of the mixed-ploidy population AG035. Colored dots indicate cytotyped individuals, colors indicate ploidy with diploids being blue, triploids purple and tetraploids orange.

**Fig. S2:**
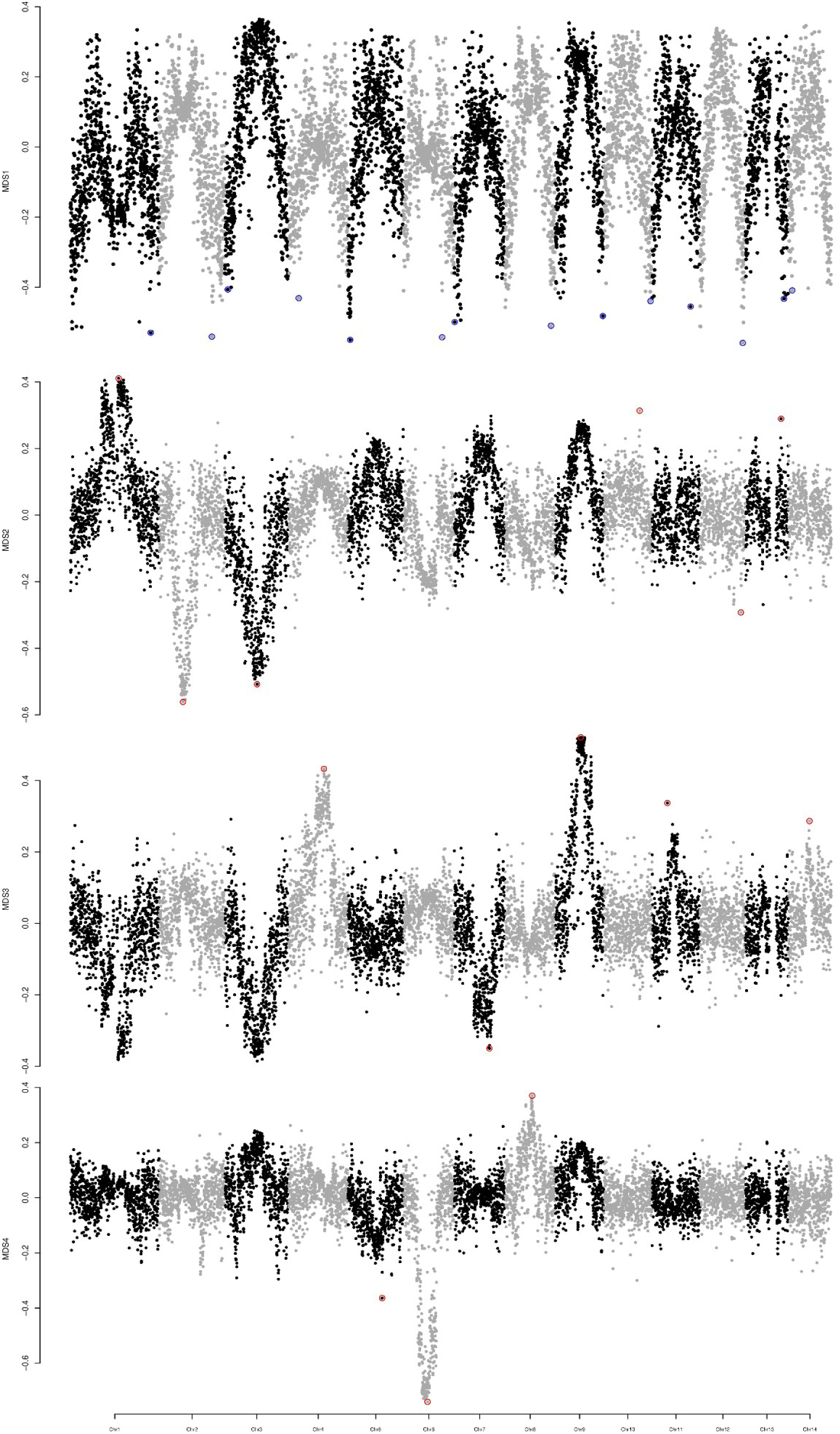
Results of the MDS analyses for PCAs in 50kb windows for contact zone AG008. Rows represent the four MDS dimensions, windows highlighted in blue present values from chromosome arms with PCAs plotted in Fig. S3, results highlighted in red indicate outliers with PCAs plotted in Fig. S4.

**Fig. S3:**
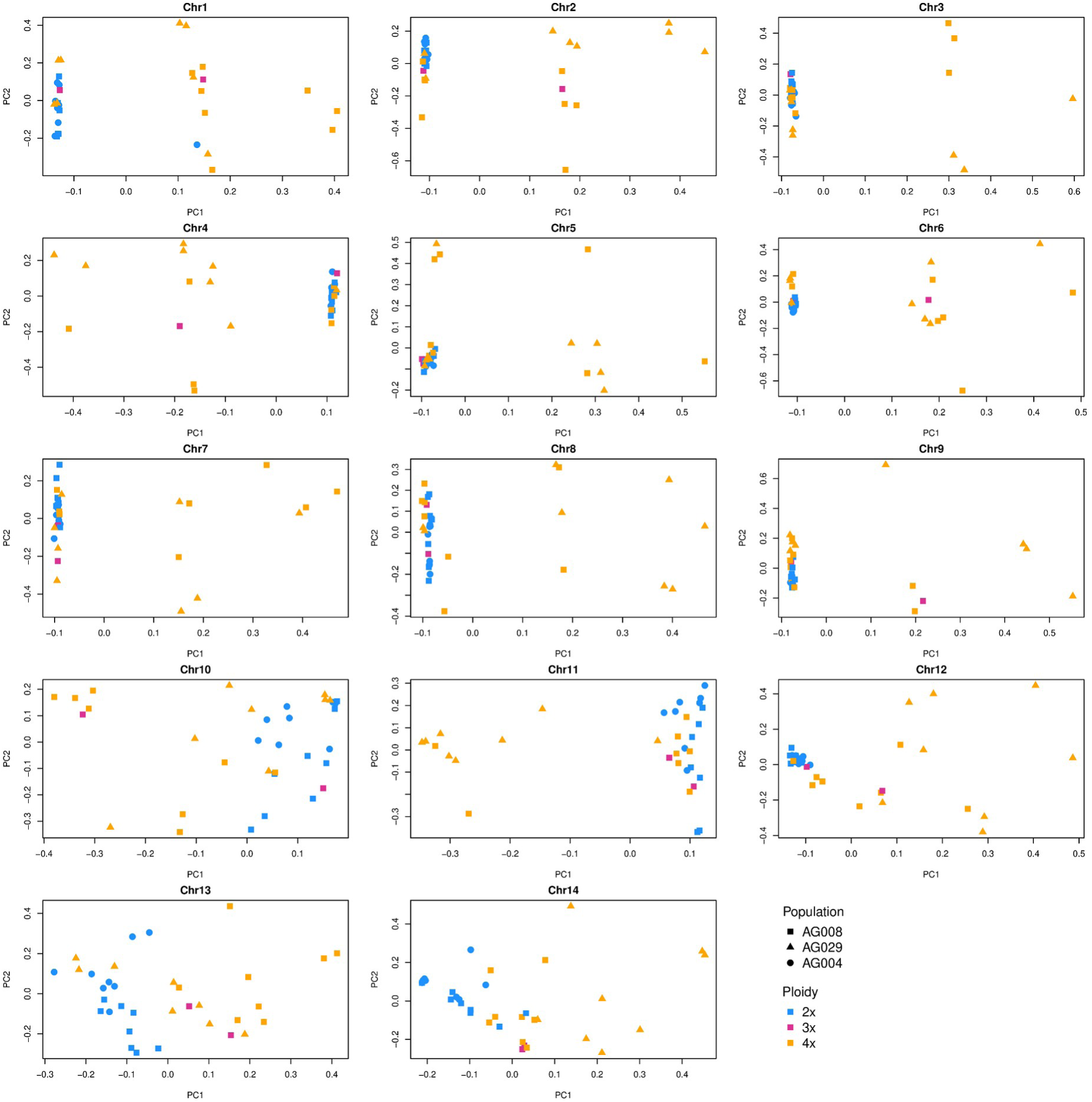
local PCAs for outlier windows for mixed-ploidy population AG008 identified using MDS. Each plot shows results from a local PCA based on an outlier window identified in Fig. S1 for each of the chromosomes.

**Fig. S4:**
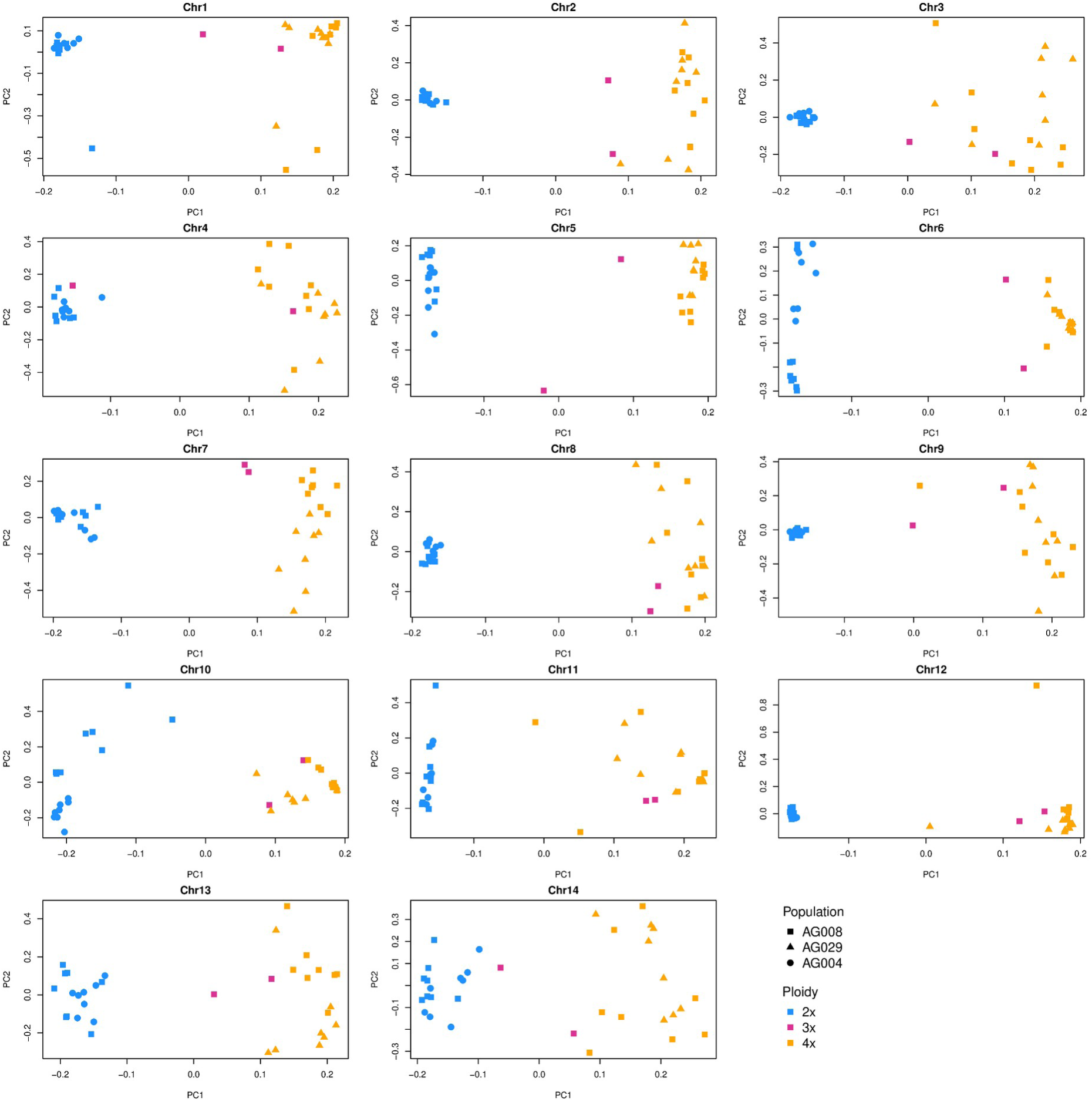
local PCAs from chromosome arms for mixed-ploidy population AG008. Each plot shows results from a local PCA from a window with lowest values for MDS1 identified in Fig. S1 for each of the chromosomes.

**Fig. S5:**
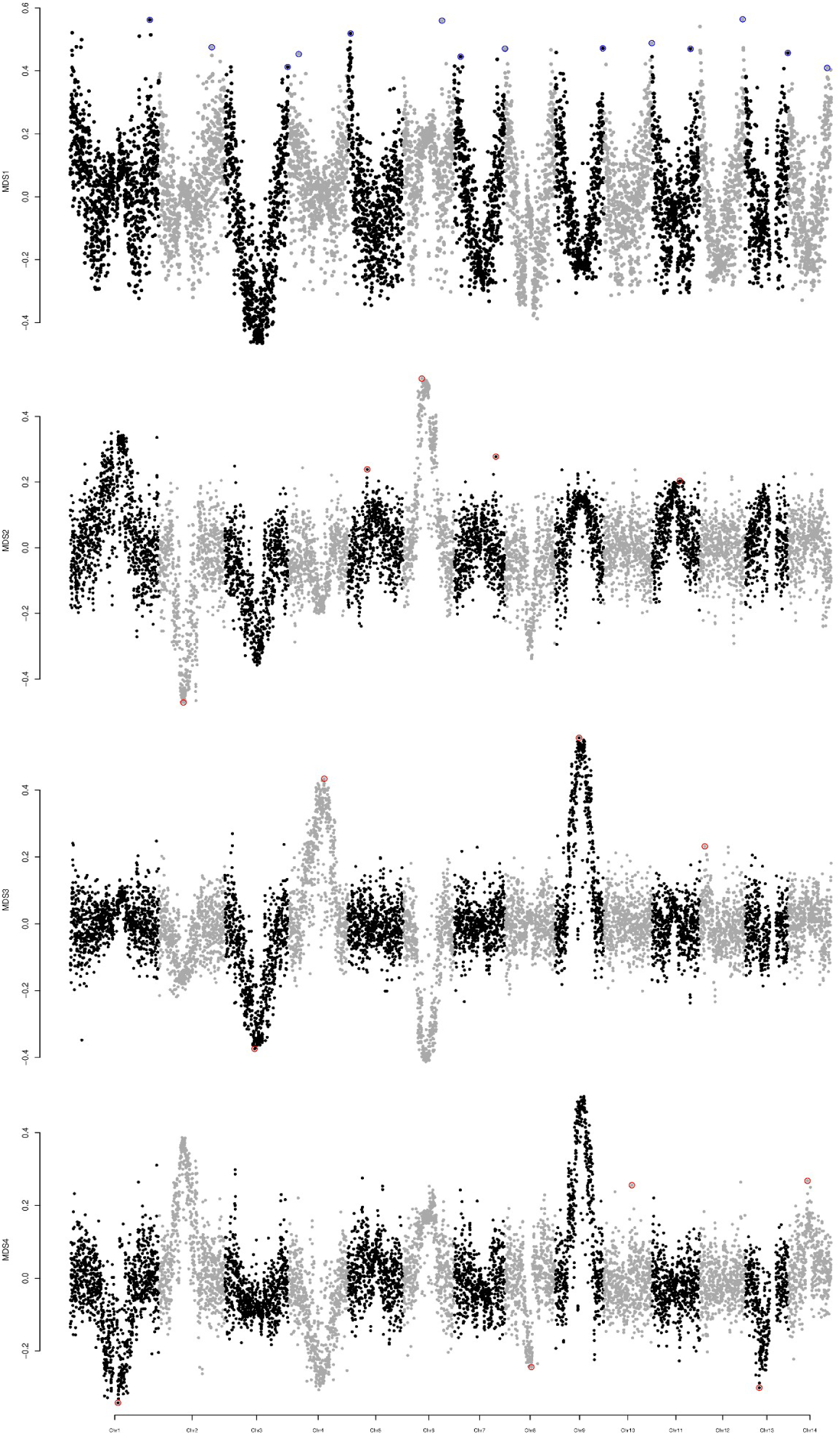
Results of the MDS analyses for PCAs in 50kb windows for contact zone AG030. Rows represent the four MDS dimensions, windows highlighted in blue present values from chromosome arms with PCAs plotted in Fig. S6, results highlighted in red indicate outliers with PCAs plotted in Fig. S7.

**Fig. S6:**
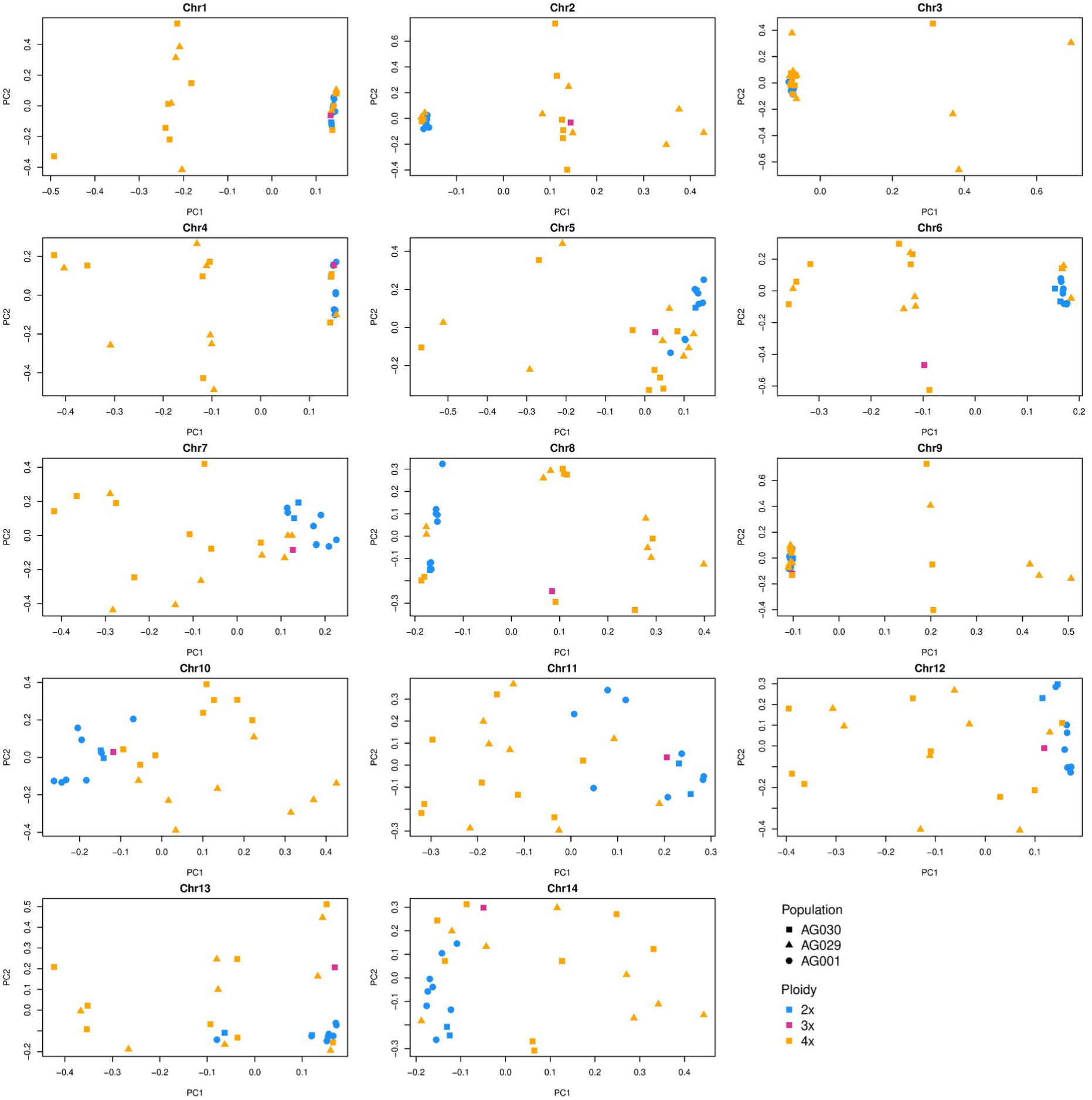
local PCAs for outlier windows for mixed-ploidy population AG030 identified using MDS. Each plot shows results from a local PCA based on an outlier window for one of the chromosomes identified in Fig. S5.

**Fig. S7:**
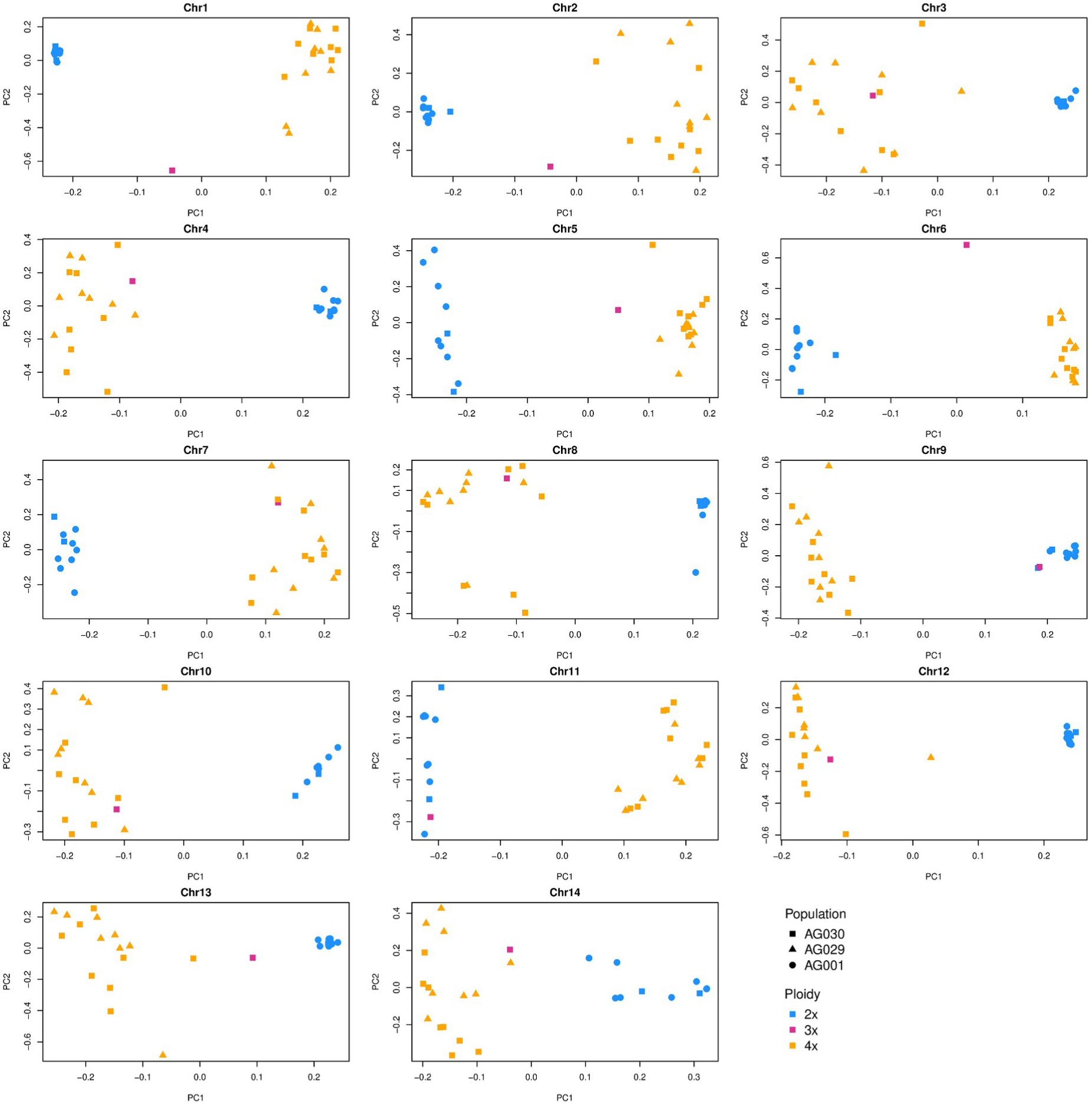
local PCAs from chromosome arms for mixed-ploidy population AG030. Each plot shows results from a local PCA from a window with highest values for MDS1 identified in Fig. S5 for each of the chromosomes.

**Fig. S8:**
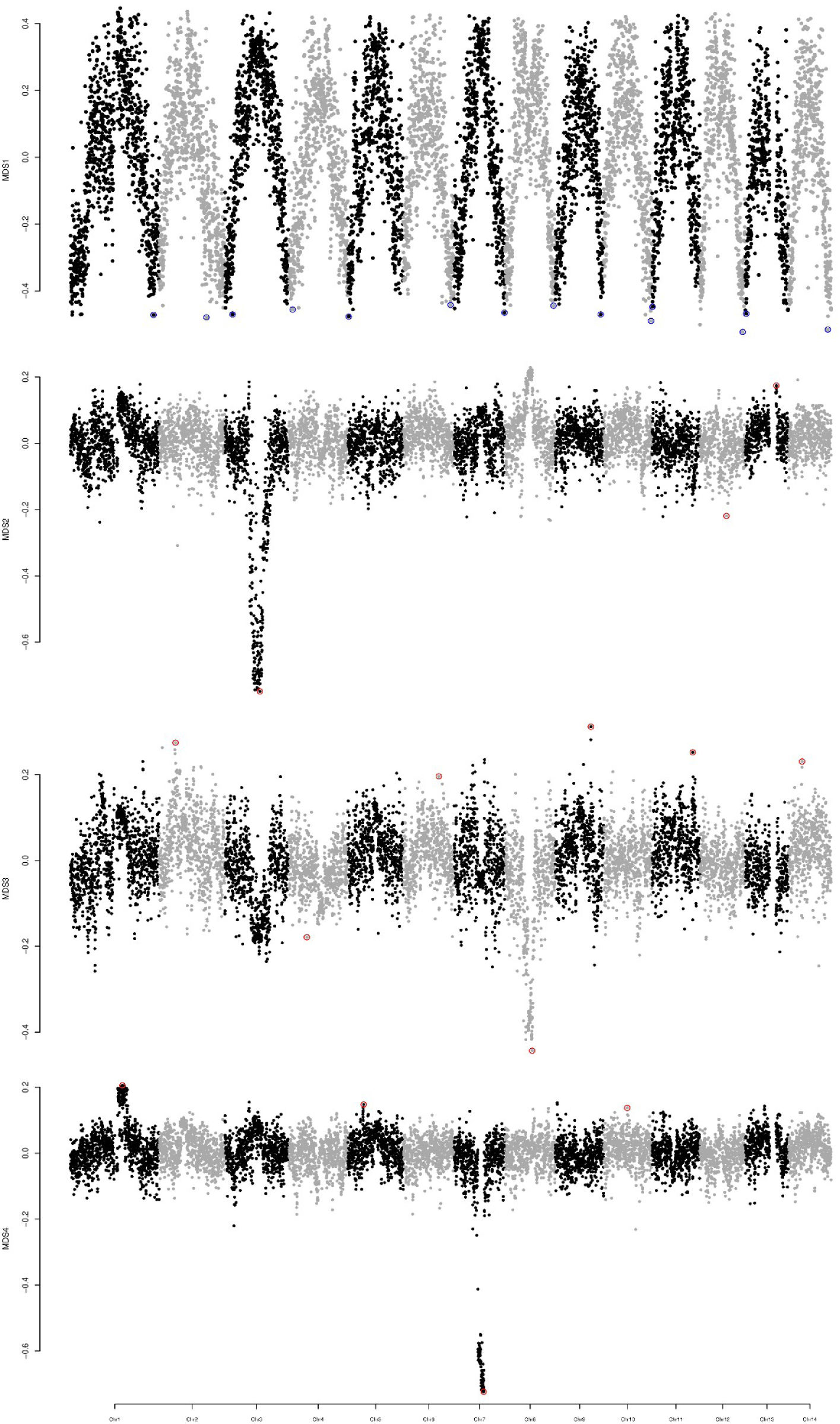
Results of the MDS analyses for PCAs in 50kb windows for contact zone AG035. Rows represent the four MDS dimensions, windows highlighted in blue present values from chromosome arms with PCAs plotted in Fig. S9, results highlighted in red indicate outliers with PCAs plotted in Fig. S10.

**Fig. S9:**
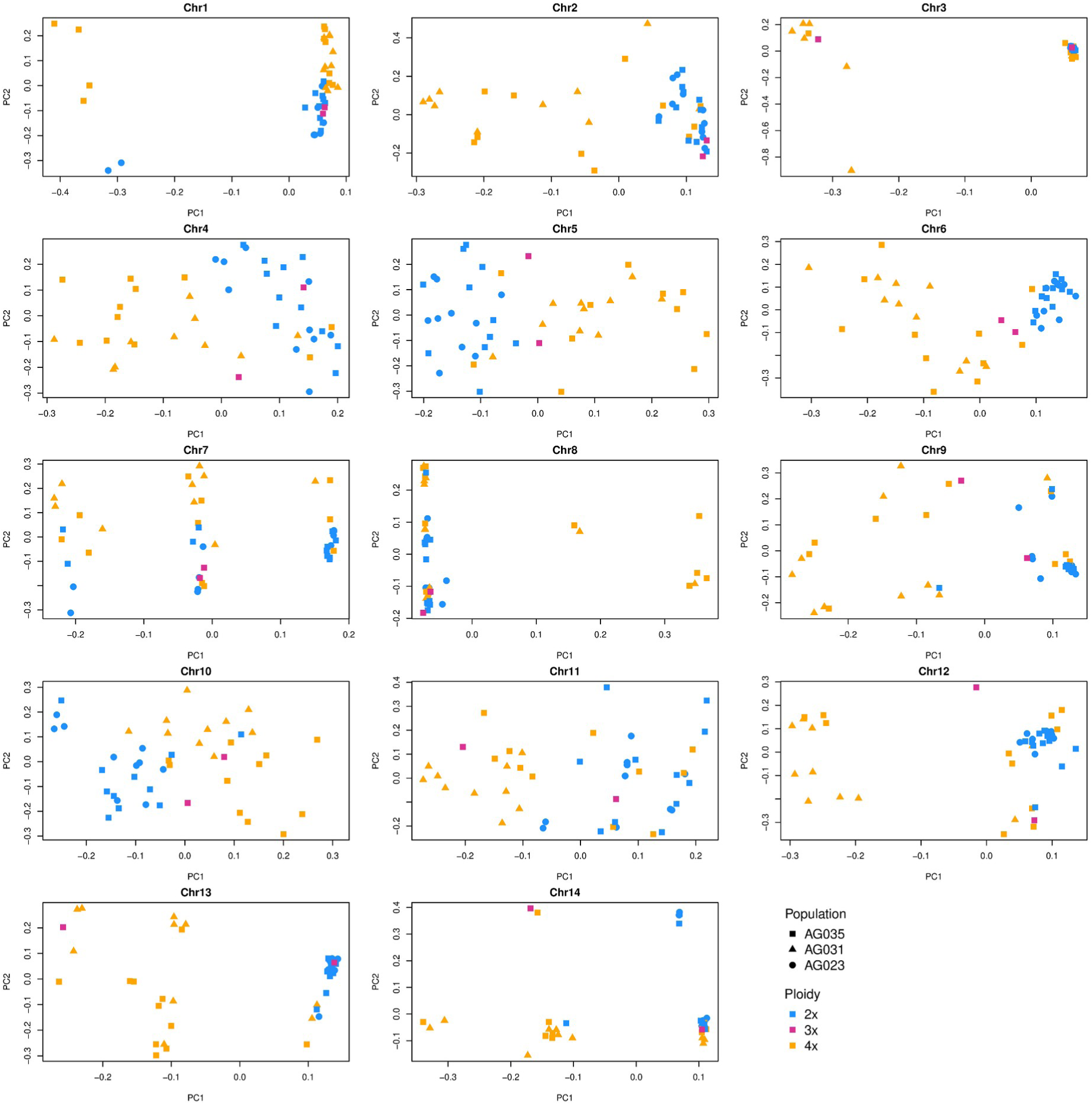
local PCAs for outlier windows for mixed-ploidy population AG035 identified using MDS. Each plot shows results from a local PCA based on an outlier window for one of the chromosomes identified in Fig. S8.

**Fig. S10:**
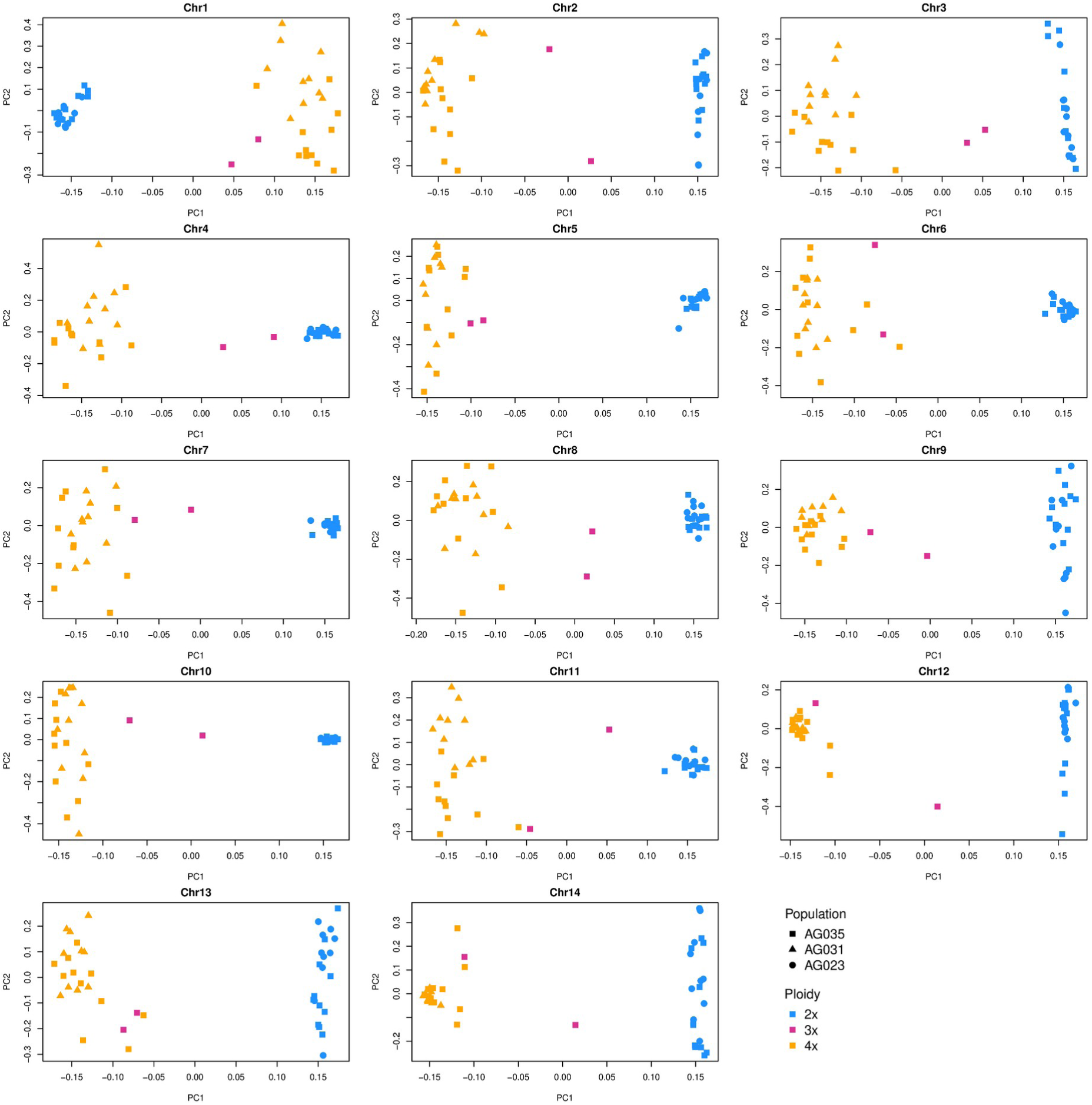
local PCAs from chromosome arms for mixed-ploidy population AG035. Each plot shows results from a local PCA from a window with lowest values for MDS1 identified in Fig. S8 for each of the chromosomes.

## Notes

### Competing Interest Statement

The authors have declared no competing interest.

